# Freshwater input and tidal position regulate species turnover and interaction rewiring in intertidal ecological networks

**DOI:** 10.64898/2026.06.10.731491

**Authors:** Anthony J. Gillis, Mads S. Thomsen, Derek Gerber, Daniel Hernandez-Carrasco, Jonathan D. Tonkin

## Abstract

The effect that environmental conditions have on community and network assembly processes remains unclear, in part because these processes operate at multiple scales. Because marine primary producers and microinvertebrates have limited mobility, are susceptible to multiple stressors, and can be observed interacting *in situ*, their habitat-based interactions provide an informative system for disentangling network organising processes. We sampled 646 habitat-use networks, quantifying interactions involving ‘habitat-users’ and biogenic ‘habitat-formers’ over 12 months at 9 sites within Te Ihutai/Avon-Heathcote estuary in Christchurch, Aotearoa New Zealand. Using generalised dissimilarity mixed-effect models, we examined whether changes to species interactions – deconstructed into species turnover and interaction rewiring – were modulated by environmental covariates, including freshwater discharge, elevation, temperature, spatial location and season. We found that with increasing dissimilarity in sites proximity to freshwater, interaction change was more driven by rewiring, whereas differences in elevation (i.e., between channels and non-channel habitats) were driven by species turnover, with more sessile species inhabiting tidal channels. The proximity of habitats also played a strong role, with nearby networks comprising more similar interactions, and species turnover becoming more prevalent with increasing distance. Our results highlight that the relative influence and magnitude of rewiring and species turnover in controlling estuarine interaction networks was affected by the individual species distributions across the estuary and their responses to separate, but co-occurring, environmental factors. Quantification of habitat-former/user interaction networks offers robust, albeit understudied, measures of processes that can underpin community assembly, highlighting their potential importance in research, management and conservation.

**Open research statement:** Data are provided for peer review. The code for produced from the project analysis and used to draft this manuscript is shared via a public GitHub repository hosted by the Tonkin Research group (repo name: *<u>EstInteractTurn</u>*). The data used in the formal analysis is hosted on Zenodo, under the corresponding authors profile (doi: 10.5281/zenodo.20619091). The data and code was prepared following strict adherence to the FAIR principles, meaning all data was saved as comma-separated values (*.csv*) or native R data structures (.*rds*).

## Introduction

Understanding how community assembly is influenced by habitat selection, species interactions, and environmental conditions is a fundamental goal of community ecology (Chase, 2003; Drake, 1990; Flores-Arguedas et al., 2023; Fukami, 2015). The assembly of and changes to species interaction networks, in particular, depends on various processes that operate at multiple spatial and temporal scales (Marjakangas et al., 2022; Ponisio et al., 2019). Disentangling these processes is critical to better understand how interactions networks vary across space and through time (Carstensen et al., 2014; Simanonok & Burkle, 2014), but requires approaches that account for changes to both species composition and interactions. Overall changes in species interactions (i.e., the total interaction turnover), can be partitioned into ‘species turnover’ (β*_st_*), and ‘interaction rewiring’ (β*_rw_*), reflecting changes in both composition and interaction partners (within a shared species pool) between networks, respectively (Fig. 1; Fründ, 2021; Novotny, 2009; Poisot et al., 2012).

**Figure 1.**
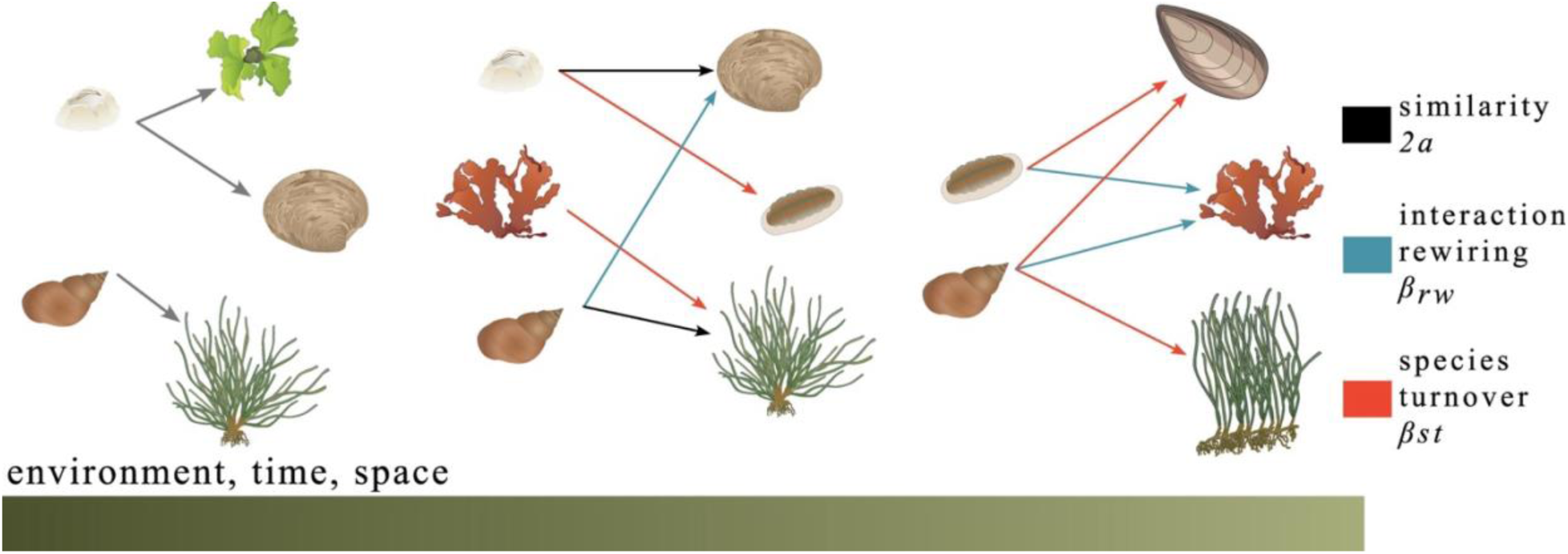
Simplified interaction networks showing habitat-users attached to habitat-formers (arrow direction) as they change along environmental gradients, through time or across spatial location (green gradient). Arrow colour represents how interactions remain the same (black), or change due to interaction rewiring (blue), or species turnover (red) compared to the previous network along the gradient.

The contribution of interaction turnover mechanisms to network assembly depends, in part, on species-specific responses to key environmental factors (Bascompte & Stouffer, 2009; Chase, 2003; Ponisio et al., 2019). Species turnover can contribute to interaction turnover when the species composition of networks changes due to niche selection or partitioning along strong deterministic environmental gradients, such as altitude (with associated differences in temperature and precipitation; Sekar et al., 2024; Simanonok & Burkle, 2014). In contrast, rewiring, i.e., partner switching among the same pool of species, can contribute to interaction turnover when the relative abundances of some species change along spatiotemporal gradients in environmental conditions, and resource availability (CaraDonna et al., 2017; Ceron et al., 2022; Lázaro & Gómez-Martínez, 2022). Given the varied species responses to environmental conditions, it remains unclear how the different turnover mechanisms contribute to network assembly, particularly in spatiotemporally dynamic systems (Pellissier et al., 2018; Ward et al., 2026).

Coastal bar-built low-energy estuaries are complex, interconnected, and highly variable habitats, formed and maintained by geophysical processes, including the deposition of sediment from rivers and the ocean, and fluctuations in sea levels. Estuaries are generally dominated by soft sediments and burrowing infauna, like worms and bivalves (Levin et al., 1996). By contrast, estuaries have little hard substrate, such as rocks and boulders, for seaweeds and ‘epifaunal’ macroinvertebrates to attach to – epibiotic life strategies that instead dominate on rocky reefs.

Nevertheless, estuarine organisms can form complex epibiotic interaction networks, where and when habitat-forming organisms, including seaweeds or cockles, provide attachment, feeding and hiding space for habitat-using organisms, such as snails and crabs (Clemente & Thomsen, 2025b; Manca et al., 2022; Thomsen et al., 2016). Furthermore, the limited mobilities of the many epibiotic seaweeds and invertebrates that make-up habitat-based interaction networks may increase their susceptibility to stress from multiple co-occurring environmental factors, potentially revealing new insights into network assembly processes (Domínguez et al., 2021; Krug et al., 2021; Wang et al., 2024). For example, the vertical tidal position (e.g., along a tidal flat-to-tidal channel gradient) may filter species that are susceptible to desiccation from networks (reflecting species turnover; Chang et al., 2018). In contrast, rewiring may become more prevalent along a freshwater-influenced salinity gradient where stenohaline snails become more abundant and use a wider variety of habitat-formers further from freshwater sources. Finally, metacommunity and spatial processes, like dispersal, may affect how functionally connected communities are, thereby modualting interaction similarity between networks (Novotny, 2009). Because disentangling the complexity of the competing processes that underpin network assembly is challenging with traditional dissimilarity-based methods, modern hierarchical approaches, such as Generalised Dissimilarity Mixed effect Models (GDMM; Hernández-Carrasco et al., 2026; White et al., 2024; Woolley et al., 2017), are required.

Here, we combined network theory and GDMMs to investigate the mechanisms underpinning assembly of interaction networks and their variability in response to the predictability of key stressors in a dynamic estuarine ecosystem. Using networks formed by intertidal habitat-former-user interactions, we assessed how the strength of species turnover and rewiring was modified by pairwise dissimilarities in distance to nearest river, tidal elevation, season, air temperature, freshwater discharge, and inter-sample distance. We hypothesised that rewiring would become more important to overall interaction turnover with increasing dissimilarity in distance to freshwater source (H_1a_) and freshwater discharge (H_1b_) because freshwater inputs may have opposing effects on different species (e.g., near-river sites with low salinity and high organic loading may support euryhaline filter-feeders but suppress stenohaline grazers). We also hypothesised that species turnover would contribute more to the overall interaction turnover along strong deterministic filtering conditions, including tidal elevation (H_2a_), temperature (H_2b_) or season (H_2c_), partly because many species are poorly adapted to desiccation stress, which is exacerbated at high air temperatures during the summer months.

Lastly, we hypothesised that networks that are close to each other in space will have relatively similar species pools and interactions (hereafter referred to ‘similarity’) and will exhibit increasing species turnover with increasing spatial distance (H_3_).

## Methods

### Study site

Ihutai/Avon-Heathcote is a ca. 7 km^2^ bar-built tidal estuary located in Ōtautahi/Christchurch in Aotearoa/New Zealand (Fig. 2a; Foster, 2019; Gerber, 2021). The estuary receives freshwater (and associated nutrients and sediments) from the northern Ōtākaro/Avon and southwestern Ōpāwaho/Heathcote rivers and fully saline water through a small inlet located in the southeast corner (Fig. 2b). These sources of fresh and marine water create a salinity gradient in the estuary from ca. 4 to 34 ppt, depending on location and temporally varying tidal and riverine flow regimes (Gerber, 2021; Jones & Simons, 1981; Marsden, 2004). Moreover, tidal channels which remain submerged during low tide, and tidal flats, which typically are emerged during low tide, create a strong desiccation gradient (Foster, 2019; Gerber, 2021; Jupp et al., 2007). The tidal amplitude varies from 1.7 to 2.2 m, exposing large tidal flats to atmospheric conditions for 3-5 hours during low tide (Foster, 2019; Gerber, 2021; McClatchie et al., 1982). In the estuary, bare sand and mud flats are interspersed with biogenic habitat-formers, like the seagrass *Zostera mulleri*, the seaweeds *Ulva* spp. and *Gracilaria chilensis*, high densities of the endemic cockle, *Austrovenus stutchburyi* (which can often protrude partly above the sediment surface to provide an attachment space for epibiota) and scattered dead shells deposited on the sediment (Clemente & Thomsen, 2024, 2025a, 2025b). *Z. muelleri*, *Ulva* spp., and *A. stutchburyi*, are archetypical estuarine foundation species that create critical habitat and attachment substrates for epiphytes (e.g., filamentous algae), epifaunal sessile species (e.g., barnacles), and epifaunal species with limited mobility (e.g., limpets; Clemente & Thomsen, 2025a, 2025b; Gerber, 2021).

**Figure 2.**
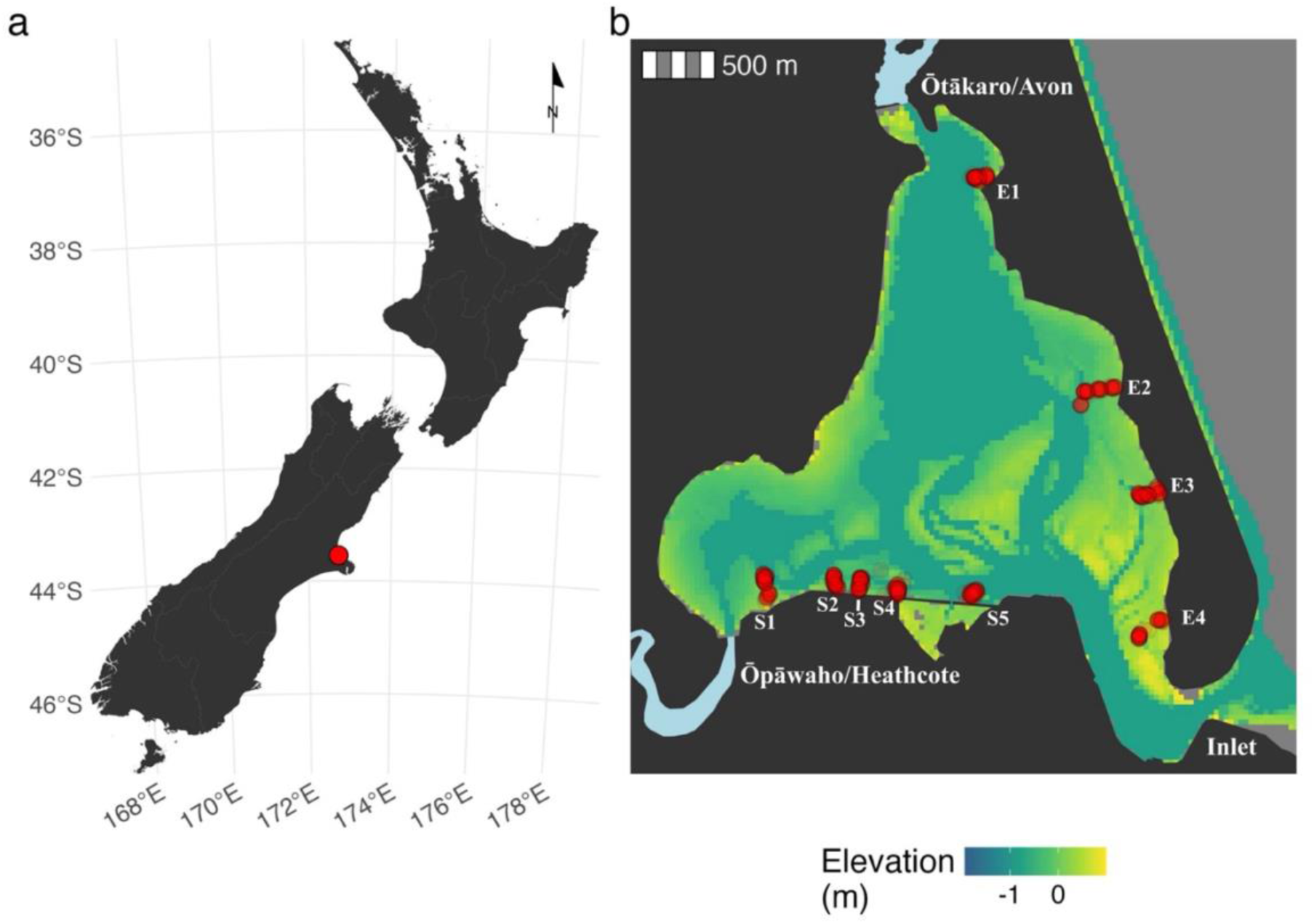
(a) Location of Ihutai/Avon-Heathcote estuary, near Ōtautahi/Christchurch, New Zealand, and (b) quadrat sampling sites (S1-5 & E1-4) where interactions were recorded (red), with two river locations that feed the estuary depicted in light blue.

### Sampling of habitat-former/user interactions

Using an established habitat-former-user interaction sampling protocol (see Montie et al., 2024; Montie & Thomsen, 2023; Thomsen & South, 2019), we recorded interactions between habitat-formers and habitat-users ca. two hours before low tides, each month from November 2019 to October 2020 (except for April 2020, where sampling was prohibited due to COVID-19 restrictions). Each month, we haphazardly placed 0.0625 m^2^ georeferenced quadrats at nine sites positioned along the estuary’s eastern and southern shores (a total of ca. 60 samples per month, all samples were separated by > 2m, Fig. 2b). To quantify the role of vertical elevation, approximately half of the quadrats were sampled on the tidal flats and half in tidal channels. Within each quadrat, we recorded the interactions between the biogenic (e.g., *Z. mulleri* and *A. stutchburyi*) and non-living (e.g., mud, dead shells) habitat-formers and their associated epibiotic habitat-users. Habitat-users could either be physically attached to the habitat-former (e.g., sessile species like barnacles or slow-moving species like limpets) or living directly underneath the habitat-formers (e.g., many snails and crabs, underneath seaweed fronds). We did not quantify infauna in the sub-surface sediments (typically polychaetes and bivalves) in this study. All species were identified, and their specific habitat interaction were recorded, such as *Elminius* (a barnacle) attached to *A*. *stutchburyi* or *Micrelenchus huttoni* (a snail) attached to *Z. muelleri*. In addition to two-species interactions, we also recorded longer interaction chains, such as *Membranipora* spp (a bryozoan) on *M. huttoni* on *Ulva* spp. on *A. stutchburyi* (an example of a rare four-level chain).

### Environmental and spatial covariates

We used the proximity to nearest freshwater sources (i.e., distances to the two river mouths) and LIDAR-derived vertical elevation (relative to mean sea level) as proxies for salinity and desiccation stress, respectively. Using the *sf* package in R (Pebesma, 2018), we calculated the minimum distance (m) to the nearest river mouth and extracted elevation (m) for each quadrat from a 1-metre resolution digital raster elevation model obtained from Land Information New Zealand (LINZ: https://data.linz.govt.nz/layer/121859-new-zealand-lidar-1m-dem/). To account for the potential role of dispersal, we also calculated the spatial distance (m) between all quadrats using a spatial distance function from the *sf* package in R (Pebesma, 2018). Furthermore, hourly air temperature (°C) data were downloaded from the Bromley weather station (agent no: 43967) using the National Institute of Water and Atmosphere (NIWA) data repository DataHub (https://data.niwa.co.nz/products/climate-station-hourly). Additionally, daily freshwater flow data (m^3^/s) were downloaded from the Environmental Canterbury (ECAN) Open Data portal (https://data.dev.ecan.govt.nz/Catalogue/Method?MethodId=79) from the Buxton Terrace and Gloucester Street Bridge flow gauges, representing freshwater flow from the Heathcote and Avon rivers, respectively. Using the downloaded freshwater and temperature data, we then calculated the mean freshwater flow and mean air temperature for the month prior to sampling to account for any prolonged time periods of high flow and temperature that may repeatedly affect the epibiota communities.

### 3.3 Statistical analysis

#### Patterns in community and interaction composition

We visualised the multivariate community for the entire estuary (pooled across time) and monthly variability (with 95% confidence ellipses) at specific sampling sites with non-metric multidimensional scaling (NMDS) ordination, using the Sørensen dissimilarity index with the *ecodist* R package (Goslee & Urban, 2007). We visualised network structure with chord diagrams by pooling network adjacency matrices at the site level and arranging them by increasing distance to the nearest freshwater river (Fig. 2b), using the *circlize* R package (Gu et al., 2014). We then used a generalised linear model to assess whether interaction count and richness varied across months, tidal position and sites, which were ordered based on their minimum proximity to a river using the function *glmmTMB* and a negative binomial distribution (*nbinom2*; M. Brooks et al., 2017). Last, we conducted an analysis of deviance with type III sums of squares and a χ^2^ test using the *car* package and the *Anova* function to assess the amount of variation explained by each factor in our glm.

### Interaction turnover, covariate distances, and GDMM

The composite of two networks can be decomposed into the contributions of (a) interaction rewiring, (b) species turnover (*st*) and (c) similarity (sim, Fig. 1; Fründ, 2021). Following Frund (2021), we quantified rewiring (*rw*), species turnover (*st*) and similarity (*sim*) between quadrats *i* and *j* using the Sørensen-Dice dissimilarity index (*D_SØR_ = (b + c)/(2a + b + c)*) as:

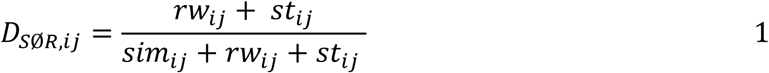

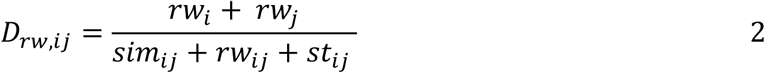

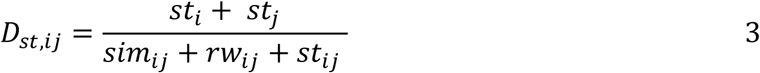

where *rw*, *st*, and *sim* are the counts of interactions attributed to rewiring, species turnover and similarity, the latter of which is denoted as *2a* in the standard index formula (Fig. 1).

To better understand the relative contribution of rewiring and species turnover to overall network changes, we modelled their contribution to network dissimilarity simultaneously using a multinomial likelihood within a generalised dissimilarity mixed model (GDMM), an extension of generalised dissimilarity models (Dias et al., 2022; Ferrier et al., 2007; Mokany et al., 2022; see Woolley et al., 2017 for use of binomial distribution). We modelled observed pairwise changes in interaction counts (*rw*, *st*, and *sim*, the numerators of Eq. 2 and 3) as coming from an underlying probability vector (θ*_ij_*), and the total number of interactions observed in both sites (*n_ij_*, the denominator of Eq. 1-3).

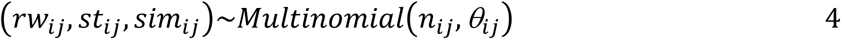

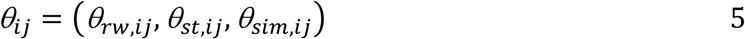

The above probability vector represent the expected proportion of interactions contributing to turnover or rewiring (θ*_rw,ij_*, θ*_st,ij_*), which is mathematically equivalent to their respective contribution the Sørensen dissimilarities ((θ*_rw,ij_* = E(*D_rw,ij_*), θ*_st,ij_* = E(*D_st,ij_*); Eq. 2 and 3). Therefore, to estimate θ*_ij_* as a function of environmental covariates (*m*), we defined our linear predictor as:

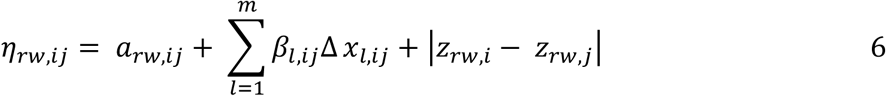

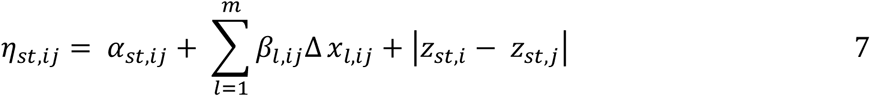

Here α is a global intercept, and β is the modelled coefficient for each covariate included in the model and Δ*x* represents the quadrat-specific pairwise environmental distance as the absolute difference in predictors (proximity to freshwater, elevation, and the previous month’s mean flow and temperature (|xi – xj|; Mokany et al., 2022; White et al., 2024). For seasonal distances, we calculated the absolute minimum difference in days between quadrat sampling dates, where distances ranged between 0 and 180 days to capture seasonal circularity. For example, two observations recorded 350 days apart should be more similar in seasonal conditions (despite their distant in time). Additionally, inter-quadrat spatial distance was included as a model-predictor to account for spatial autocorrelation and the role of spatial processes, like dispersal. Lastly, we addressed non-independence of pairwise observation by incorporating a latent variable *z* denoting sample-level (here, quadrat-level) errors into the model structure, and included the absolute pairwise differences in the linear predictor (*|z_i_ - z_j_|*; White et al., 2024).

We mapped our linear predictors (η) for each mechanism to the probability vector (θ*_ij_*) using the *softmax* function:

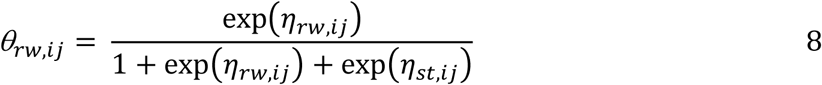

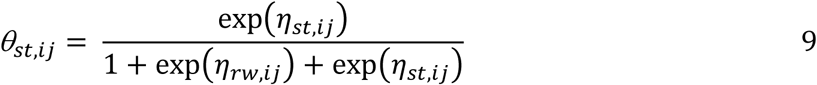

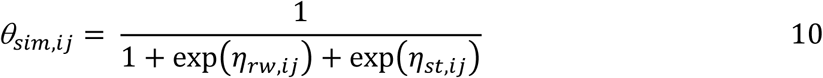

Interaction similarity (θ*_sim,ij_*) was designated as our reference category in the multinomial distribution because our hypotheses are focused on the comparative relationship between species turnover and rewiring. Before model fitting, we centred and scaled covariate distance and used the Nelder-Mead optimiser to estimate initial parameter values, which improved chain convergence (Schielzeth, 2010; White et al., 2024). Using the *greta* package in R (Golding, 2019), we then fit the model with a Bayesian Monte Carlo Markov chain sampler (MCMC), consisting of 4 chains and leapfrog step size set to 25 (min) and 30 (max) to help with efficient sampling. We selected to use uninformative priors for the intercepts and environmental coefficients, such that α *∼ N*(*0, 0.5*) and β *∼ N*(*0, 0.5*). We used regularising priors to alleviate computational pressures associated with hyper-prior distributions, such as *z ∼* N(*0, 1*) × σ and σ *∼ half-N*(*0, 0.1*). Each chain was warmed up with 8000 iterations and sampled for 5000 iterations. We visually inspected posterior distributions and calculated the Gelman and Rubin potential scale reduction factor to ensure model chains converged (*R* < 1.1; Brooks & Gelman, 1998; Gelman & Rubin, 1992; Vehtari et al., 2021).

## Results

### Patterns in community composition and network interactions

We sampled a total of 646 quadrats which included 28 living epibiota taxa (i.e., seaweeds, seagrasses, and macroinvertebrates) and 2 types of non-living substrates (dead shells and mud). Visual inspection of the NMDS plot, suggested that both proximity to freshwater and vertical elevation influenced community composition (Fig. 3a). However, temporal and site-level variability contributed to substantial overlaps in the 95% confidence ellipses on the NMDS plot (Fig. 3b).

**Figure 3.**
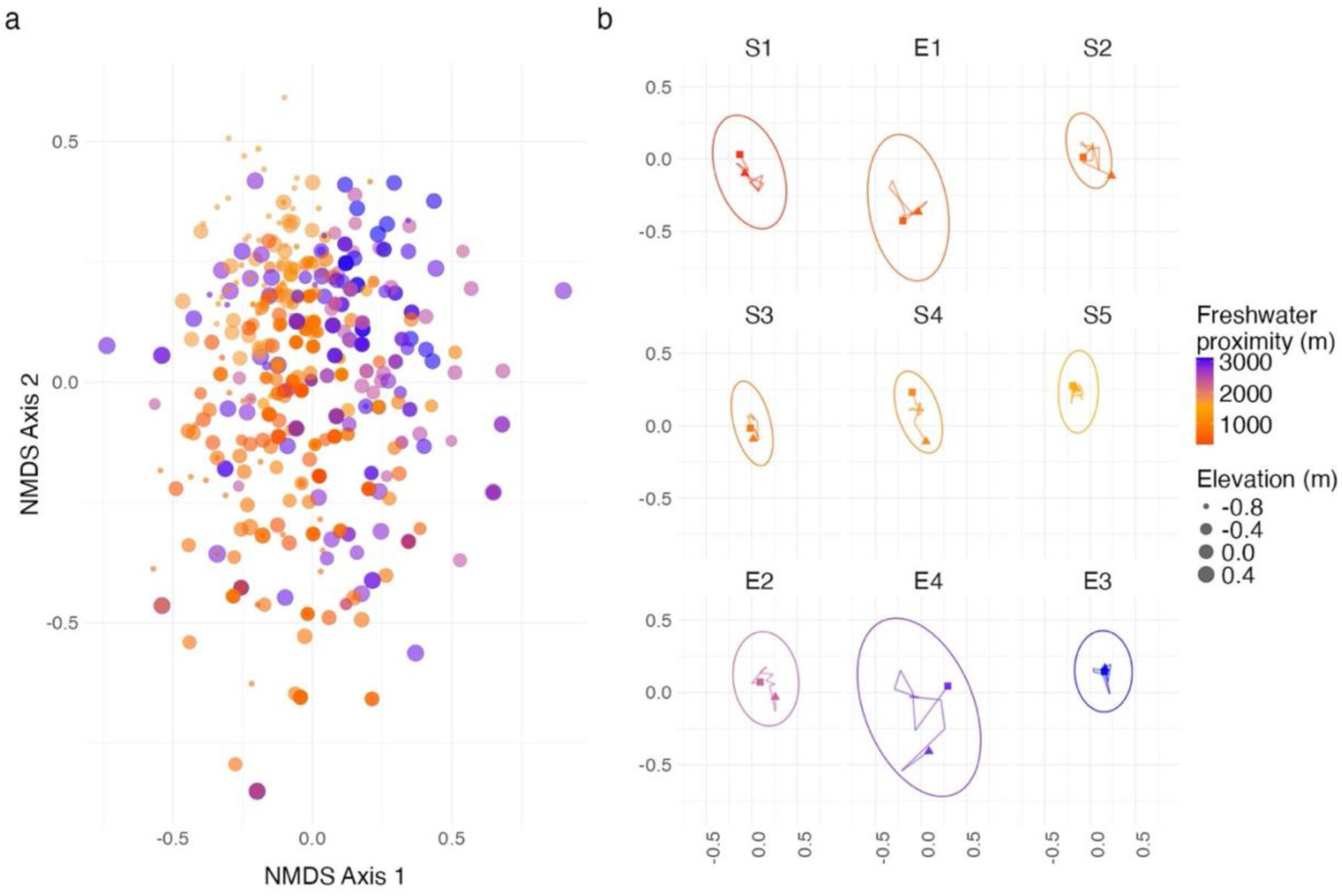
(a) Non-metric multidimensional scaling ordination (NMDS, *n* = 464, stress = 0.24, r^2^ = 0.79) representing community composition in the Ihutai/Avon-Heathcote estuary, where point-size and point-colour reflects vertical elevation and freshwater proximity, respectively. (b) Site-level NMDS plots depicting monthly changes in community centroids between November 2019 (square) to October 2020 (triangle). The total number of quadrats (*n*) sampled at each site were: S1 = 77, E1 = 76, S2 = 65, S3 = 77, S4 = 74, S5 = 73, E2 = 73, E4 = 53 and E3 = 78.

Across the year of sampling, we observed a total of 127 unique interactions involving the 30 habitat-formers and habitat-users. We found a significant interaction between the variance explained by site and tidal position on interaction count and interaction richness (count – *p* = 0.5e^-11^, χ^2^ = 69.6; richness – *p* = 0.007, χ^2^ = 20.8). Broadly, we observed an increase in interaction counts and richness towards the middle of the gradient and at lower tidal positions (see sites S2-5; Fig. 4a, c). Furthermore, we found a significant amount of variation in interaction count and richness explained by month (count – *p* = 0.1e^-17^, χ^2^ = 108.5; richness – *p* = 0.2e^-14^, χ^2^ = 91.7), where both measures were highest during winter (Jun-Aug) and early- to mid-spring (Sep, Oct) at lower tidal elevations (Fig. 4b, d). The most important habitat-formers included dead shells, mud, and *Z. muelleri* (but this habitat-former had a limited distribution along the eastern coastline only; Fig. 5). Moreover, *Diloma subrostrata, M. huttoni* and *Ulva* spp. were observed many times as both habitat-formers and habitat-users, interacting with multiple partners. Finally, *M. huttoni*, and *Notoacmea helmsi* were amongst the most numerous habitat-users. Examining our networks revealed that closer proximity to freshwater led to networks dominated by a greater incidence of interactions between *A. stutchburyi*, *D. subrostrata*., *Ulva* spp., and mud. Network and interaction composition became more diverse with increasing distance to the nearest freshwater source, partly because *N. helmsii*., *M. huttonii,* and *Z. muelleri* became more abundant. Dead shell became a more important habitat former further away from freshwater sources where the number of interactions with *A. stutchburyi* decreased (Fig. 5).

**Figure 4.**
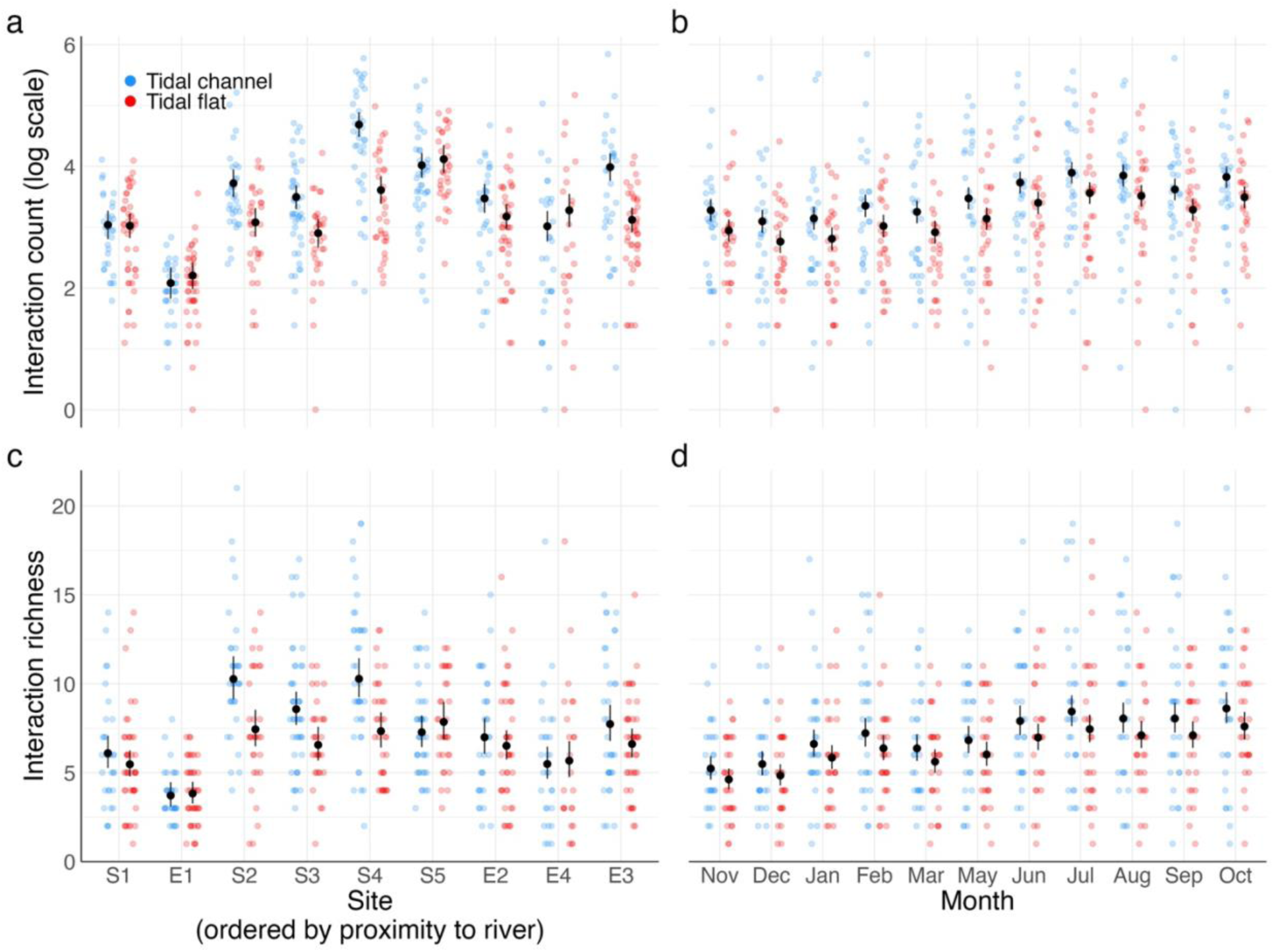
(a, b) The total number of interactions counted and (c, d) interaction richness (the number of unique interactions) observed in each network (quadrat size = 0.0625 cm^2^) across sites (ordered by distance to nearest freshwater source), month (beginning in 2019 and ending in 2020) and tidal position. Black points represent the GLM estimated marginal means, and the black lines the 95% confidence intervals.

**Figure 5.**
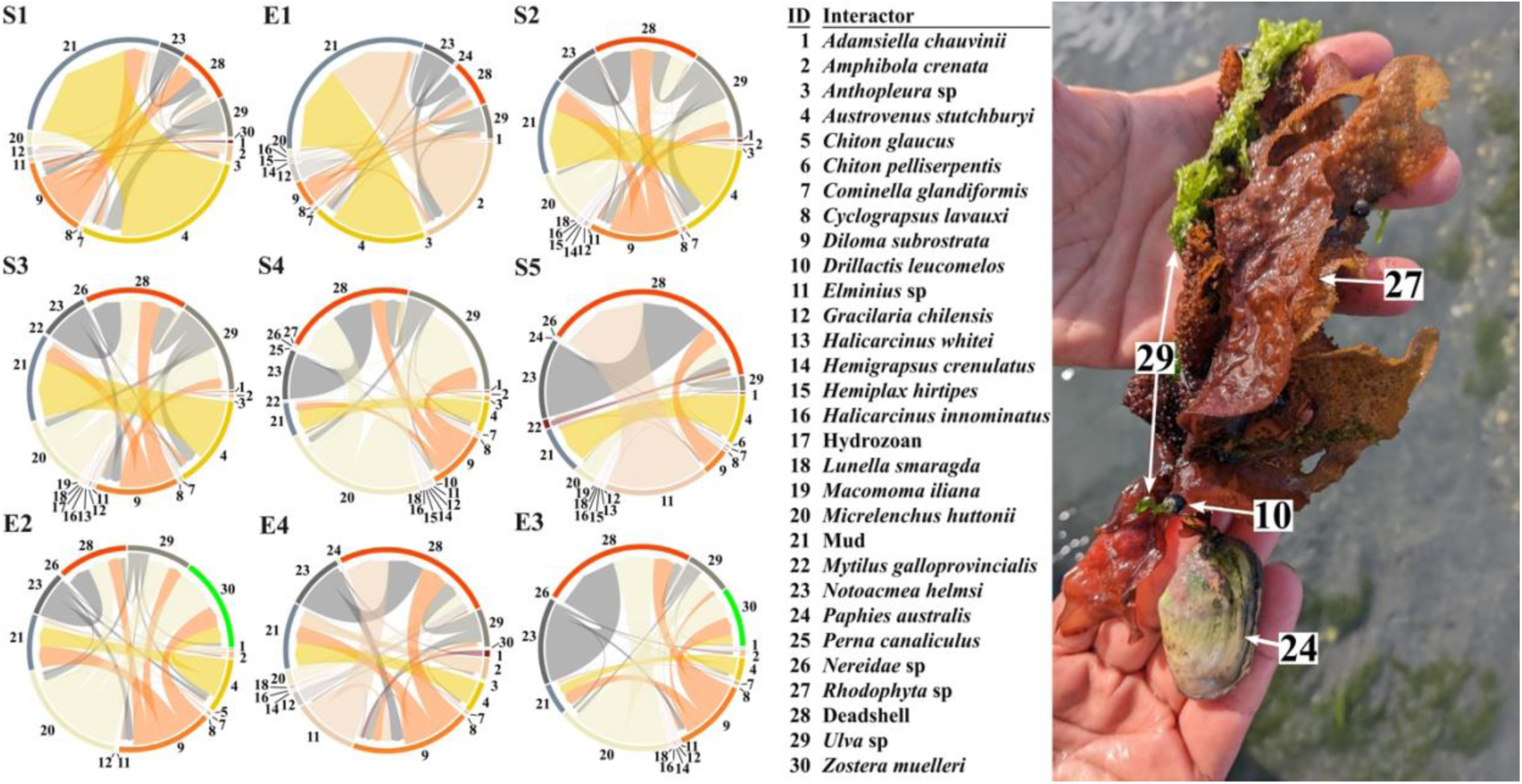
Attachment interaction networks across nine sites in the Avon-Heathcote estuary, ordered according to distance from the nearest freshwater source. The directional arrows depict habitat users interacting with habitat formers. Link colours were classified by the habitat-user to separate different species. Link width was determined by the number of observations (n) between habitat-users and habitat-formers at each site (S1 = 77, E1 = 76, S2 = 65, S3 = 77, S4 = 74, S5 = 73, E2 = 73, E4 = 53 and E3 = 78). The inset photo (credit: Anthony J Gillis) represent a visual example of an attachment network between macroinvertebrates (10, 24) and seaweeds (27, 29).

### Interaction turnover, GDMMs

We found that with increasing difference in distance to nearest freshwater source, interaction similarity and species turnover decreased, while the contribution of rewiring to overall interaction turnover increased (Fig. 6a). Furthermore, species turnover increased with increasing elevation and temperature distances, while rewiring and interaction similarity decreased (Fig. 6b and c). By contrast, increasing distances in temperature, seasonality (days) and freshwater flow showed small increases in species turnover, and decreases in rewiring and similarity (Fig. 6c-e). Finally, we found that increasing spatial distance resulted in the greatest changes in network structure, where species turnover contributed more to the overall interaction change (Fig. 6f). Conversely, when networks were in close proximity differences between rewiring, species turnover and similarity were marginal (Fig. 6f). When diagnosing our models, we found that our model successfully converged as all *R* point estimate remained below 1.1.

**Figure 6.**
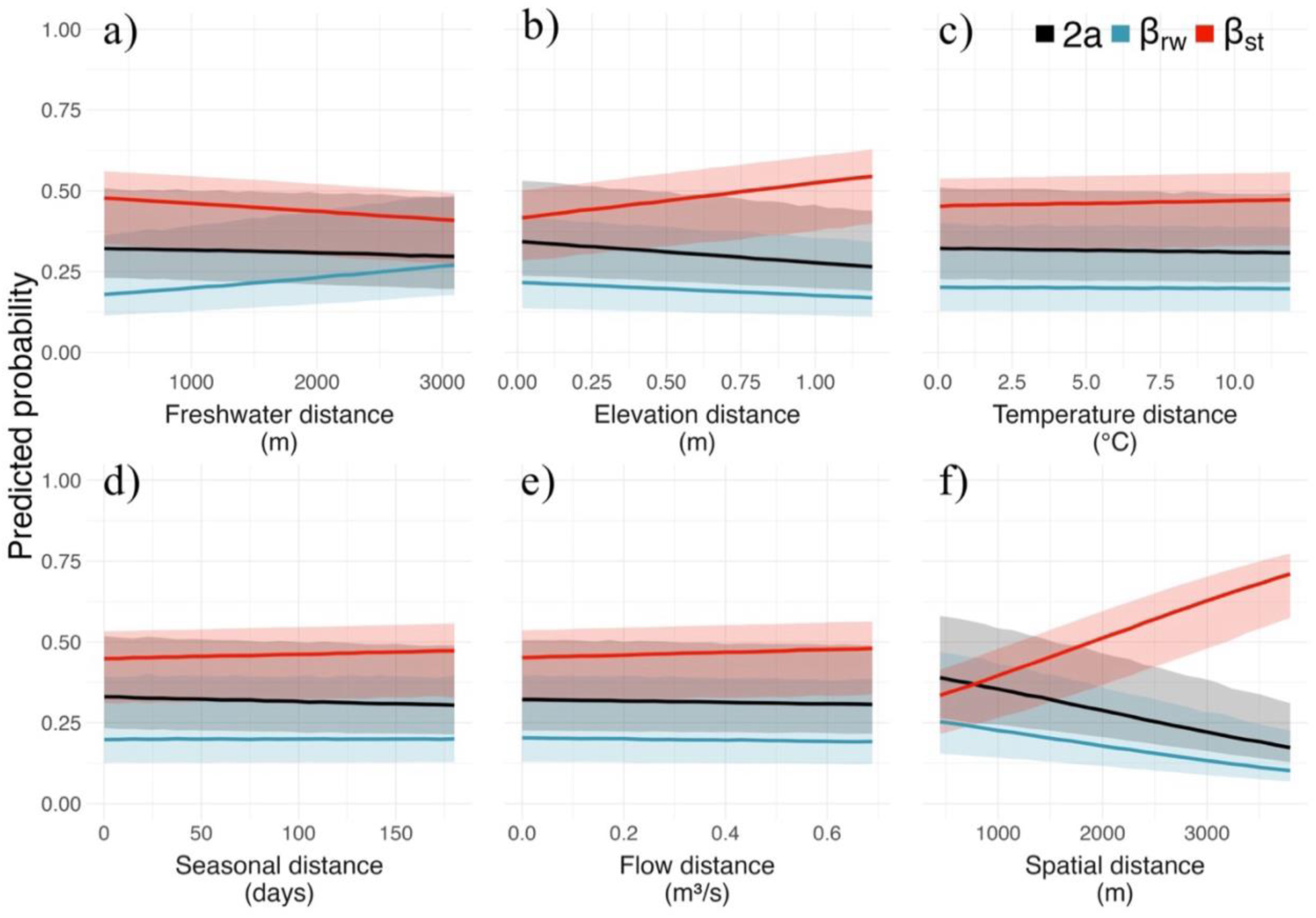
The mean probability and 95% credibility thresholds (shaded areas) of observing the different interaction turnover mechanism (blue – rewiring, β_rw_, and red – species turnover, β_st_) or similar interactions (similarity – 2a) across (a) proximity to freshwater, (b) elevation, (c) seasonal, (d) spatial, (e) temperature and (f) flow distance gradient in the Ihutai/Avon-Heathcoat estuary.

## Discussion

Our results demonstrated that the processes underpinning network assembly in a dynamic estuary differed depending on several factors, including which species were filtered from habitats by co-occurring environmental stressors. Specifically, as networks became more dissimilar in their proximity to freshwater, the probability of interaction rewiring increased, driven, in part, by reciprocal occurrences of specific habitat-forming species (supporting H_1a_). Conversely, species turnover increased with increasing distance in tidal elevation, likely driven by strong environmental filtering of desiccation-sensitive species, limiting them to lower elevations (supporting H_2a_). Furthermore, networks that were closer together were more likely to have similar interactions compared to distant networks, where the overall interaction change was driven by species turnover (supporting H_3_). This latter result suggests that dispersal is limited between locations in the estuary. Finally, using a relatively novel joint modelling method with a multinomial distribution enabled us to identify the relative contributions of the different turnover mechanisms that underpin network assembly in dynamic ecosystems, and how these mechanisms vary in strength with co-occurring environmental stress factors.

### Freshwater input, interaction richness and rewiring

Proximity to freshwater source, which alters salinity, nutrients and sedimentation stressors, impacted network structure through increased rewiring, driven, in part, by the reciprocal dominance of key habitat-forming species. For example, *A*. *stutchburyi*, an important filter-feeding habitat-former, was relatively more common at sites closer to the two rivers, potentially facilitated by high levels of suspended organic matter and lower salinities, compared to the most marine sites (Marsden, 2004; Sandwell et al., 2009). In contrast, dead shells and photosynthesising habitat-formers such as *Ulva* spp. and *Z. muelleri* were observed more frequently at sites farther from the rivers, where suspended sediments was likely lowest and water clarity highest (Clemente & Thomsen, 2025a; Siciliano et al., 2019). In response to these shifting habitat-formers, relatively stress tolerant habitat-users, such as *D. subrostrata* or *N. helmsi*, likely selectively associated with the locally dominant habitat-formers. However, stenohaline habitat-users, such as *M. huttoni,* also contributed to the increase in rewiring, as they were observed more frequently in networks closer to the marine inlet and interacted with a greater diversity of habitat-formers.

As a result of these reciprocal species responses and sustained by the positive habitat feedback mechanisms (Bulleri et al., 2016), the sites in between rivers and the ocean had the greatest overlaps in interacting species distributions and therefore supported a high number of total and unique species interactions (Roshni et al., 2022; Sharpe & Baldwin, 2009). This mid-domain spatial overlap in species niches and increased rewiring mirrors patterns observed in subalpine plant-pollinator networks during mid-season, when a higher phenological overlaps occur in flowering plant species and pollinators that diversify their interaction partners (CaraDonna et al., 2017).

### Tidal position and elevation as a strong community filter

In contrast to the reciprocal species responses to differences in freshwater proximity, vertical tidal position limited several species from higher elevation thereby resulting in high species turnover among tidal positions. Mirroring the stressful conditions associated with systems along altitudinal gradients, where extreme temperature and humidity are typically limiting factors (Kraft et al., 2011; Leahy et al., 2024; Simanonok & Burkle, 2014; Sponsler et al., 2022), high elevational tidal flats comprised low interaction counts and interaction richness. More specifically, we found that many sessile species, including barnacles, bristle worms, and hydrozoans, which cannot mitigate heat and desiccation stresses by moving to damp microclimates, were largely absent from networks on the tidal flats (Fig. S1). These results were unsurprising as desiccation and warming are archetypal stressors known to affect several marine species, limiting their distributions to lower tidal positions (Boese et al., 2005; Harley, 2011) and thus altering community structure between tidal elevations (Amstutz et al., 2021; Hillebrand et al., 2010).

### Spatial distance and dispersal limitations guide metacommunity organisation

Our analysis also demonstrated that interaction similarity was largest when networks were closer together in space. Gastropods, such as *A*. *crenulata*, *D*. *subrostrata* and *M*. *huttoni*, which move relatively short distances to connect close-proximity habitats (Darcy & Eggleston, 2005; Gillanders et al., 2003), likely played a key role in maintaining interactions in nearby communities. Additionally, elevated freshwater input, during floods or cyclical tidal inundation likely promoted interaction similarities via momentary passive dispersal events (Jiajun et al., 2024). However, these forms of habitat-connectivity were relatively limited (∼ 700 m, see Fig. 6f), as species turnover increasingly contributed to overall interaction turnover at greater spatial distances, suggesting that dispersal limitation likely influenced the formation of these networks (Forbes et al., 2025; Novotny, 2009; Padial et al., 2014).

### Temporal and seasonal variability

Temporal changes and seasonality only had minor effects on structuring networks in the Avon-Heathcote estuary, as reflected by the negligible responses of different turnover mechanisms to seasonal distance. The lack of seasonal turnover may have been driven by the high abundance of relatively long-lived macroinvertebrates and clonal seagrasses that buffer seasonal fluctuations, (Clemente et al., 2023; Curtis et al., 2000), compared to a fewer number of ephemeral or seasonal algae species in the communities (Clemente & Thomsen, 2024, 2025a; Gauna et al., 2017). Indeed, the monthly variability in interaction counts may have been driven more by short-term fluctuations in species-specific abundances, such as following local storms or anomalous river flows, which do not translate to seasonal changes in network structure. Similarly, community composition did not exhibit clear or consistent temporal changes either within or across sites, and monthly variability was minor (except for site E4). Perhaps the lack of seasonal changes in these estuarine networks could be expected, as New Zealand more broadly is considered to have relatively muted seasonality in many environmental conditions (such as rainfall and temperature), due to its maritime-influenced climate (Hernández-Carrasco et al., 2025; Tonkin et al., 2017, 2018).

### Future research directions

Here we highlighted the importance of sessile life history traits in modulating habitat-use interactions in estuaries results from their strong dependence on lower elevation habitats. Clearly, other functional and morphological traits affect species distribution in estuaries, and thus the pool of potential interaction partners, and, even if species can physically interact (i.e., trait matching; Dehling et al., 2014; Peralta et al., 2020; Tylianakis & Morris, 2017). For example, the size of intertidal species, such as gastropods and bivalves, has been tied to thermal tolerance, energy expenditure, and survival, which can limit their distributions among microhabitats (Peck et al., 2009). We generally observed more mobile species in similar proportions across tidal elevations (Fig. S1), but smaller individuals of mobile molluscs, such as *A. crenulata*, may have indirectly contributed to species turnover due to their limited capacity to provide attachment space on their shells. Future studies of seaweed-macroinvertebrate networks could benefit from measuring and analysing traits known to modify species interactions, including their sizes, feeding modes, life-histories, dispersal capabilities and chemical deterrents (Boström et al., 2010; Pereira & Da Gama, 2008; Veríssimo et al., 2024).

The increasing probability of rewiring across freshwater input gradients suggests a potential role of widespread interaction generalism (high diversity of interaction partners) throughout the estuary. Typically, aquatic macrophyte-animal networks have a relatively high proportion of interaction generalism, which can contribute to the persistence of these networks (Bates & DeWreede, 2007; Manca et al., 2022; Taylor & Cole, 1994). However, several studies have shown that aquatic macrophyte-epifauna networks are often structured by specialist interactions between specific sets of partners (Manca et al., 2025; Montie et al., 2024; Thomsen & South, 2019). Indeed, our observation of high species turnover suggests that the networks in the Avon-Heathcote may also contain more specialist interactions. Still, there is a limited number of analyses on coastal habitat-based interactions (Montie et al., 2024; Montie & Thomsen, 2023; Thomsen & South, 2019), and more studies should therefore quantify how interaction generalist and specialist taxa contribute to the turnover and persistence of networks in dynamic coastal ecosystems.

Finally, we note that this analysis is a first joint modelling approach for quantifying species turnover, rewiring and interaction similarity using a multinomial distribution with GDMMs. In using a multinomial distribution, we could quantify the magnitude, direction of change, and uncertainty of each category’s response to environmental variability while accounting for the response of the other categories. This analytical approach allowed us to more accurately model and better assess each category’s contribution to real-world interaction similarities and turnover along key environmental gradients (CaraDonna et al., 2017; Woolley et al., 2017). Still, by using interaction dissimilarities derived from binary presence/absence metrics, we may have underestimated the contributions of turnover mechanisms compared to dissimilarity components derived from interaction counts (e.g., using link weights). The use of interaction count-derived dissimilarities could potentially have revealed a greater contribution of rewiring along the proximity to freshwater gradients or demonstrated seasonal trends (cf. Fig. 5). Further development of GDMMs could incorporate and test network turnover using more quantitative approaches, such as combing weighted interaction counts with the total abundances of both habitat-formers and -users (CaraDonna et al., 2017; Fründ, 2021; Simanonok & Burkle, 2014).

## Conclusion

Our study revealed that co-occurring environmental factors can have varied effects on interacting estuarine species, shaping their distributions and ultimately affecting the processes that modulate network assembly. Differences in proximity to rivers led to reciprocal abundance responses in key habitat-forming species, often resulting in interaction changes among habitat users and dominant habitat forming species. By contrast, tidal position and elevation contributed more to species turnover often due sessile species being limited to lower, damper tidal channels. Dispersal limitation also likely influenced the assembly of these estuarine networks, with a clear influence on species turnover with increasing distance. Ultimately, our study showed that, in the Avon-Heathcote estuary with many co-occurring environmental stressors, the estuarine networks were structured by a few habitat-forming species that facilitated local biodiversity (Clemente & Thomsen, 2025b).

Our findings were supported by applying novel multinomial GDMMs, which jointly modelled the relative contribution of interaction turnover mechanisms and enabled us to more accurately analyse and disentangle the combined contributions of rewiring and species turnover to network assembly (CaraDonna et al., 2017). New research that embeds species traits and disentangles the prevalence of interaction generalist or specialist species will help move the field forward, particularly when coupled with advanced modelling frameworks like those used here.

Analysis of turnover in these habitat-use networks provides valuable information on the processes that underpin network assembly and support coastal biodiversity, and how intensifying environmental variability, including from climate change, may alter these processes (Schleuning et al., 2016; van Dijk et al., 2015; Wernberg et al., 2024; Woodward, 2010).

## Acknowledgements

The authors acknowledge the Freshwater Ecology Research Group and Marine Ecology Research Group for their support as soundboards, providing feedback on the development of this manuscript.

## Funding Acknowledgements

Anthony J Gillis was supported by a University of Canterbury Doctoral Scholarship, provided by Jonathan D Tonkin through a Rutherford Discovery Fellowship, administered by the Royal Society Te Apārangi (RDF-18-UOC-007). Jonathan D Tonkin also acknowledges funding from Bioprotection Aotearoa and Te Pūnaha Matatini, both Centres of Research Excellence funded by the Tertiary Education Commission, New Zealand. Mads S Thomsen was supported by the New Zealand Ministry of Business, Innovation and Employment (Toka ākau toitu Kaitiakitanga—building a sustainable future for coastal reef ecosystems).

## Author Contributions

Anthony J Gillis conceptualised, prepared and analysed the data, and wrote the original draft manuscript under the co-supervision of Mads Thomsen and Jonathan Tonkin. Derek Gerber, with support from Mads Thomsen, designed the sampling methods and collected the interaction data used for the formal analysis. Daniel Hernandez-Carrasco aided in the development, refinement and code review of the model used in the formal analysis. All authors reviewed and provided edits for the manuscript.

## Supplemental Information

**Figure S1.**
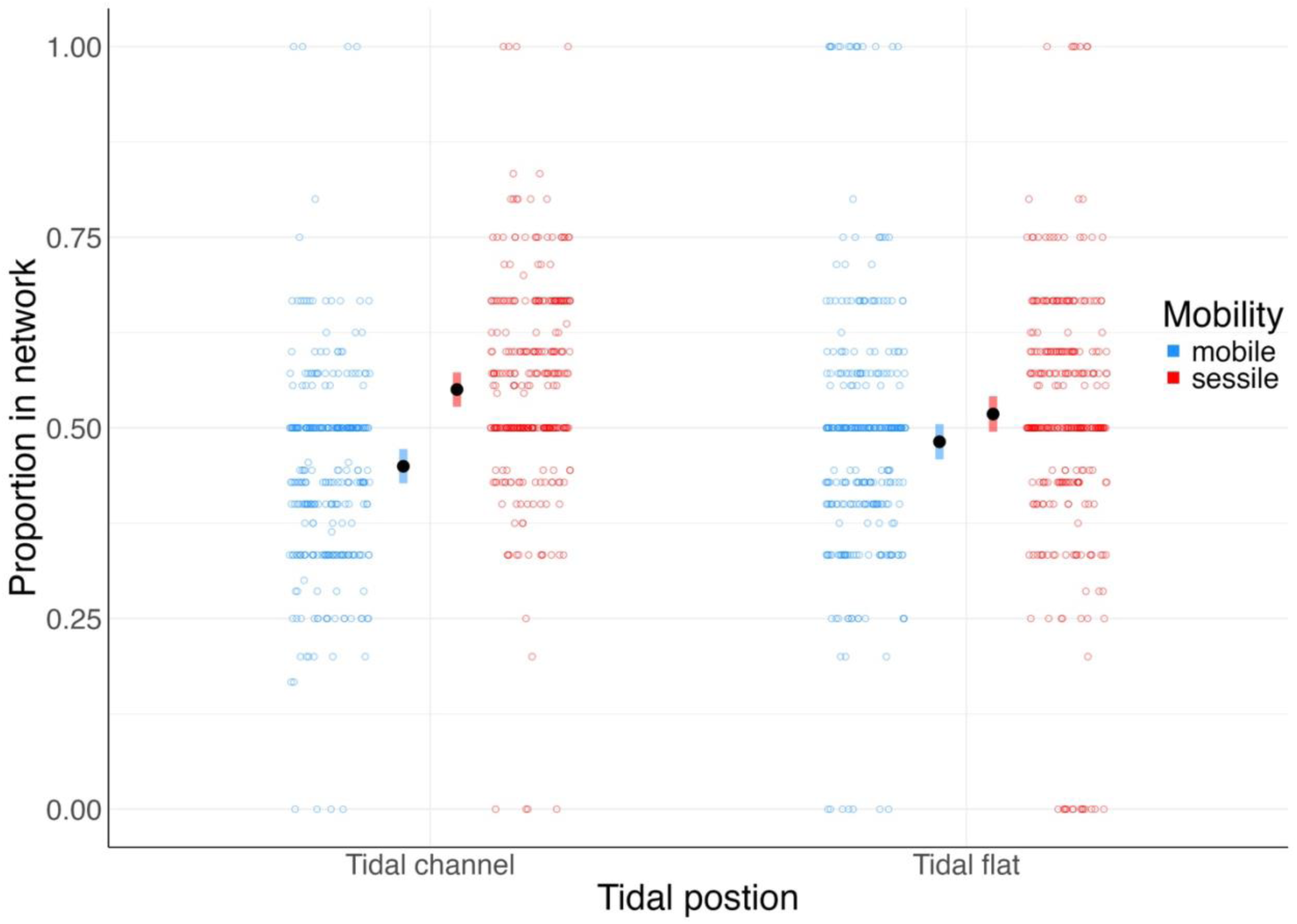
The proportion of sessile (red) or mobile (blue) species observed in seaweed-macroinvertebrate interaction networks on tidal flats and in tidal channels in the Avon-Heathcote estuary. The black bars estimated marginal means from a GLMM (with a binomial distribution) assessing the effect of tidal position (*p* = 0.051), and the coloured bars represent the 95% confidence intervals.

## References

Amstutz, A., Firth, L. B., Spicer, J. I., & Hanley, M. E. (2021). Facing up to climate change: Community composition varies with aspect and surface temperature in the rocky intertidal. Marine Environmental Research, 172. Scopus. 10.1016/j.marenvres.2021.105482

Bascompte, J., & Stouffer, D. B. (2009). The assembly and disassembly of ecological networks. Philosophical Transactions of the Royal Society B: Biological Sciences, 364(1524), 1781–1787. 10.1098/rstb.2008.0226

Bates, C. R., & DeWreede, R. E. (2007). Do changes in seaweed biodiversity influence associated invertebrate epifauna? Journal of Experimental Marine Biology and Ecology, 344(2), 206–214. 10.1016/j.jembe.2007.01.002

Boese, B. L., Robbins, B. D., & Thursby, G. (2005). Desiccation is a limiting factor for eelgrass (*Zostera marina* L.) distribution in the intertidal zone of a northeastern Pacific (USA) estuary. Botanica Marina, 48(4). 10.1515/BOT.2005.037

Boström, C., Törnroos, A., & Bonsdorff, E. (2010). Invertebrate dispersal and habitat heterogeneity: Expression of biological traits in a seagrass landscape. Journal of Experimental Marine Biology and Ecology, 390(2), 106–117. 10.1016/j.jembe.2010.05.008

Brooks, M., Bolker, B., Kristensen, K., Maechler, M., Magnusson, A., Skaug, H., Nielsen, A., Berg, C., & Van Bentham, K. (2017). glmmTMB: Generalized Linear Mixed Models using Template Model Builder (p. 1.1.13) [Dataset]. 10.32614/CRAN.package.glmmTMB

Brooks, S. P., & Gelman, A. (1998). General methods for monitoring convergence of iterative simulations. Journal of Computational and Graphical Statistics, 7(4), 434–455. 10.1080/10618600.1998.10474787

Bulleri, F., John F. Bruno, Silliman, B. R., & Stachowicz, J. J. (2016). Facilitation and the niche: Implications for coexistence, range shifts and ecosystem functioning. Functional Ecology, 30(1), 70–78. 10.1111/1365-2435.12528

CaraDonna, P. J., Petry, W. K., Brennan, R. M., Cunningham, J. L., Bronstein, J. L., Waser, N. M., & Sanders, N. J. (2017). Interaction rewiring and the rapid turnover of plant–pollinator networks. Ecology Letters, 20(3), 385–394. 10.1111/ele.12740

Carstensen, D. W., Sabatino, M., Trøjelsgaard, K., & Morellato, L. P. C. (2014). Beta diversity of plant-pollinator networks and the spatial turnover of pairwise interactions. PLoS ONE, 9(11), e112903. 10.1371/journal.pone.0112903

Ceron, K., Provete, D. B., Pires, M. M., Araujo, A. C., Blüthgen, N., & Santana, D. J. (2022). Differences in prey availability across space and time lead to interaction rewiring and reshape a predator–prey metaweb. Ecology, 103(8). 10.1002/ecy.3716

Chang, A. L., Brown, C. W., Crooks, J. A., & Ruiz, G. M. (2018). Dry and wet periods drive rapid shifts in community assembly in an estuarine ecosystem. Global Change Biology, 24(2). 10.1111/gcb.13972

Chase, J. M. (2003). Community assembly: When should history matter? Oecologia, 136(4), 489–498. 10.1007/s00442-003-1311-7

Clemente, K. J. E., & Thomsen, M. S. (2024). Facilitation cascades across space: Monitoring estuarine foundation species from satellites to the microscope. Ecosphere, 15(4), e4834. 10.1002/ecs2.4834

Clemente, K. J. E., & Thomsen, M. S. (2025a). Co-occurring foundation species increase habitat heterogeneity across estuarine intertidal environments on the South Island of New Zealand. Marine Environmental Research, 208, 107150. 10.1016/j.marenvres.2025.107150

Clemente, K. J. E., & Thomsen, M. S. (2025b). Ranking ecological contingencies from high-order factorial data demonstrate tidy control of biodiversity from facilitation cascades in estuaries on the South Island of New Zealand. Ecography, 2025(6), e07488. 10.1111/ecog.07488

Clemente, K. J. E., Thomsen, M. S., & Zimmerman, R. C. (2023). The vulnerability and resilience of seagrass ecosystems to marine heatwaves in new zealand: A remote sensing analysis of seascape metrics using PLANETSCOPE imagery. Remote Sensing in Ecology and Conservation, 9(6), 803–819. 10.1002/rse2.343

Curtis, L. A., Kinley, J. L., & Tanner, N. L. (2000). Longevity of oversized individuals: Growth, parasitism, and history in an estuarine snail population. Journal of the Marine Biological Association of the United Kingdom, 80(5), 811–820. 10.1017/S0025315400002782

Darcy, M. C., & Eggleston, D. B. (2005). Do Habitat Corridors Influence Animal Dispersal and Colonization in Estuarine Systems? Landscape Ecology, 20(7), 841–855. 10.1007/s10980-005-3704-y

Dehling, D. M., Töpfer, T., Schaefer, H. M., Jordano, P., Böhning-Gaese, K., & Schleuning, M. (2014). Functional relationships beyond species richness patterns: Trait matching in plant–bird mutualisms across scales. Global Ecology and Biogeography, 23(10), 1085–1093. 10.1111/geb.12193

Dias, F. S., Betancourt, M., Rodríguez-González, P. M., & Borda-de-Água, L. (2022). BetaBayes—A bayesian approach for comparing ecological communities. Diversity, 14(10), 858. 10.3390/d14100858

Domínguez, R., Vázquez, E., Smallegange, I. M., Woodin, S. A., Wethey, D. S., Peteiro, L. G., & Olabarria, C. (2021). Predation risk increases in estuarine bivalves stressed by low salinity. Marine Biology, 168(8), 132. 10.1007/s00227-021-03942-8

Drake, J. A. (1990). Communities as assembled structures: Do rules govern pattern? Trends in Ecology & Evolution, 5(5), 159–164. 10.1016/0169-5347(90)90223-Z

Ferrier, S., Manion, G., Elith, J., & Richardson, K. (2007). Using generalized dissimilarity modelling to analyse and predict patterns of beta diversity in regional biodiversity assessment. Diversity and Distributions, 13(3), 252–264. 10.1111/j.1472-4642.2007.00341.x

Flores-Arguedas, H., Antolin-Camarena, O., Saavedra, S., & Angulo, M. T. (2023). Assembly archetypes in ecological communities. Journal of The Royal Society Interface, 20(208). 10.1098/rsif.2023.0349

Forbes, C., Hessing-Lewis, M., & O’Connor, M. I. (2025). An Experimental Test of Dispersal Limitation of Species Diversity in Eelgrass Meadows. Estuaries and Coasts, 48(2), 35. 10.1007/s12237-024-01472-2

Foster, T. H. (2019). Distributions and interactions of co-occurring estuarine foundation species: Seagrasses, seaweeds, and shell-forming organisms. [PhD, University of Canterbury]. 10.26021/6774

Fründ, J. (2021). Dissimilarity of species interaction networks: How to partition rewiring and species turnover components. Ecosphere, 12(7). 10.1002/ecs2.3653

Fukami, T. (2015). Historical contingency in community assembly: Integrating niches, species pools, and priority effects. Annual Review of Ecology, Evolution, and Systematics, 46(1), 1–23. 10.1146/annurev-ecolsys-110411-160340

Gauna, M. C., Escobar, J. F., Odorisio, M., Cáceres, E. J., & Parodi, E. R. (2017). Spatial and temporal variation in algal epiphyte distribution on *Ulva* sp. (Ulvales, Chlorophyta) from northern Patagonia in Argentina. Phycologia, 56(2), 125–135. 10.2216/16-51.1

Gelman, A., & Rubin, D. B. (1992). Inference from iterative simulation using multiple sequences. Statistical Science, 7(4). 10.1214/ss/1177011136

Gerber, D. (2021). Associations between estuarine foundation species and their effect on biodiversity across latitudes and seasons [Master of Science]. University of Canterbury.

Gillanders, B., Able, K., Brown, J., Eggleston, D., & Sheridan, P. (2003). Evidence of connectivity between juvenile and adult habitats for mobile marine fauna: An important component of nurseries. Marine Ecology Progress Series, 247, 281–295. 10.3354/meps247281

Golding, N. (2019). greta: Simple and scalable statistical modelling in R. Journal of Open Source Software, 4(40), 1601. 10.21105/joss.01601

Goslee, S. C., & Urban, D. L. (2007). The **ecodist** package for dissimilarity-based analysis of ecological data. Journal of Statistical Software, 22(7). 10.18637/jss.v022.i07

Gu, Z., Gu, L., Eils, R., Schlesner, M., & Brors, B. (2014). *Circlize* implements and enhances circular visualization in R. Bioinformatics, 30(19), 2811–2812. 10.1093/bioinformatics/btu393

Harley, C. D. G. (2011). Climate change, keystone predation, and biodiveristy loss. Science, 334(6059), 1124–1127. 10.1126/science.1210199

Hernández-Carrasco, D., Gillis, A. J., Lai, H. R., Siqueira, T., & Tonkin, J. D. (2026). Accounting for the Influence of Community Turnover Along Environmental Gradients on Compositional Uniqueness. Ecology Letters, 29(2), e70338. 10.1111/ele.70338

Hernández-Carrasco, D., Tylianakis, J. M., Lytle, D. A., & Tonkin, J. D. (2025). Ecological and evolutionary consequences of changing seasonality. Science, 388(6750), eads4880. 10.1126/science.ads4880

Hillebrand, H., Soininen, J., & Snoeijs, P. (2010). Warming leads to higher species turnover in a coastal ecosystem. Global Change Biology, 16(4), 1181–1193. 10.1111/j.1365-2486.2009.02045.x

Jiajun, L., Biao, Z., Guangshuai, Z., Sihui, S., Yansong, L., Jinhui, Z., Jiuliang, W., & Xiangyu, G. (2024). Flooding promotes the coalescence of microbial community in estuarine habitats. Marine Environmental Research, 202, 106735. 10.1016/j.marenvres.2024.106735

Jones, M. B., & Simons, M. J. (1981). Habitat preferences of two estuarine burrowing crabs *Helice crassa* dana (grapsidae) and *Macrophthalmus hirtipes* (Jacquinot) (Ocypodidae). Journal of Experimental Marine Biology and Ecology, 56(1), 49–62. 10.1016/0022-0981(81)90007-1

Jupp, K. L., Partridge, T. R., Hart, D. E., & Marsden, I. D. (2007). Ecology of the Avon-Heathcote estuary: Comparative salt marsh survey 2006-2007 (Estuarine Research Report No. 34). Avon-Heathcote Estuary Ihutai Trust. http://hdl.handle.net/10092/1552

Kraft, N. J. B., Comita, L. S., Chase, J. M., Sanders, N. J., Swenson, N. G., Crist, T. O., Stegen, J. C., Vellend, M., Boyle, B., Anderson, M. J., Cornell, H. V., Davies, K. F., Freestone, A. L., Inouye, B. D., Harrison, S. P., & Myers, J. A. (2011). Disentangling the drivers of β diversity along latitudinal and elevational gradients. Science, 333(6050), 1755–1758. 10.1126/science.1208584

Krug, P. J., Shimer, E., & Rodriguez, V. A. (2021). Differential tolerance and seasonal adaptation to temperature and salinity stress at a dynamic range boundary between estuarine gastropods. The Biological Bulletin, 241(1), 105–122. 10.1086/715845

Lázaro, A., & Gómez-Martínez, C. (2022). Habitat loss increases seasonal interaction rewiring in plant–pollinator networks. Functional Ecology, 36(10), 2673–2684. 10.1111/1365-2435.14160

Leahy, L., Scheffers, B. R., Andersen, A. N., & Williams, S. E. (2024). Rates of species turnover across elevation vary with vertical stratum in rainforest ant assemblages. Ecography, 2024(5), e06972. 10.1111/ecog.06972

Levin, L., Talley, D., & Thayer, G. (1996). Succession of macrobenthos in a created salt marsh. Marine Ecology Progress Series, 141, 67–82. 10.3354/meps141067

Manca, F., Mulà, C., Gustafsson, C., Mauri, A., Roslin, T., Thomas, D. N., Benedetti-Cecchi, L., Norkko, A., & Strona, G. (2022). Unveiling the complexity and ecological function of aquatic macrophyte–animal networks in coastal ecosystems. Biological Reviews, 97(4), 1306–1324. 10.1111/brv.12842

Manca, F., Norkko, A. M., Cabeza, M., Llewelyn, J., Hervías-Parejo, S., & Strona, G. (2025). Macrophyte-epifauna associations form structured ecological networks in the Baltic Sea. Marine Ecology Progress Series, 771, 1–14. 10.3354/meps14942

Marjakangas, E., Muñoz, G., Turney, S., Albrecht, J., Neuschulz, E. L., Schleuning, M., & Lessard, J. (2022). Trait-based inference of ecological network assembly: A conceptual framework and methodological toolbox. Ecological Monographs, 92(2), e1502. 10.1002/ecm.1502

Marsden, I. (2004). Effects of reduced salinity and seston availability on growth of the New Zealand little-neck clam *Austrovenus stutchburyi*. Marine Ecology Progress Series, 266, 157–171. 10.3354/meps266157

McClatchie, S., Juniper, S. K., & Knox, G. A. (1982). Structure of a mudflat diatom community in the Avon-Heathcote estuary, New Zealand. New Zealand Journal of Marine and Freshwater Research, 16(3–4), 299–309. 10.1080/00288330.1982.9515973

Mokany, K., Ware, C., Woolley, S. N. C., Ferrier, S., & Fitzpatrick, M. C. (2022). A working guide to harnessing generalized dissimilarity modelling for biodiversity analysis and conservation assessment. Global Ecology and Biogeography, 31(4), 802–821. 10.1111/geb.13459

Montie, S., Schiel, D. R., & Thomsen, M. S. (2024). Shifts in foundation species dominance and altered interaction networks after compounding seismic uplift and extreme marine heatwaves. Marine Environmental Research, 202, 106738. 10.1016/j.marenvres.2024.106738

Montie, S., & Thomsen, M. S. (2023). Long-term community shifts driven by local extinction of an iconic foundation species following an extreme marine heatwave. Ecology and Evolution, 13(6), e10235. 10.1002/ece3.10235

Novotny, V. (2009). Beta diversity of plant-insect food webs in tropical forests: A conceptual framework. Insect Conservation and Diversity, 2(1), 5–9. 10.1111/j.1752-4598.2008.00035.x

Padial, A. A., Ceschin, F., Declerck, S. A. J., De Meester, L., Bonecker, C. C., Lansac-Tôha, F. A., Rodrigues, L., Rodrigues, L. C., Train, S., Velho, L. F. M., & Bini, L. M. (2014). Dispersal ability determines the role of environmental, spatial and temporal drivers of metacommunity structure. PLoS ONE, 9(10), e111227. 10.1371/journal.pone.0111227

Pebesma, E. (2018). Simple features for R: Standardized support for spatial vector data. The R Journal, 10(1), 439. 10.32614/RJ-2018-009

Peck, L. S., Clark, M. S., Morley, S. A., Massey, A., & Rossetti, H. (2009). Animal temperature limits and ecological relevance: Effects of size, activity and rates of change. Functional Ecology, 23(2), 248–256. 10.1111/j.1365-2435.2008.01537.x

Pellissier, L., Albouy, C., Bascompte, J., Farwig, N., Graham, C., Loreau, M., Maglianesi, M. A., Melián, C. J., Pitteloud, C., Roslin, T., Rohr, R., Saavedra, S., Thuiller, W., Woodward, G., Zimmermann, N. E., & Gravel, D. (2018). Comparing species interaction networks along environmental gradients. Biological Reviews, 93(2), 785–800. 10.1111/brv.12366

Peralta, G., Vázquez, D. P., Chacoff, N. P., Lomáscolo, S. B., Perry, G. L. W., & Tylianakis, J. M. (2020). Trait matching and phenological overlap increase the spatio-temporal stability and functionality of plant–pollinator interactions. Ecology Letters, 23(7), 1107–1116. 10.1111/ele.13510

Pereira, R. C., & Da Gama, B. A. P. (2008). Macroalgal Chemical Defenses and Their Roles in Structuring Tropical Marine Communities. In C. D. Amsler (Ed.), Algal Chemical Ecology (pp. 25–55). Springer Berlin Heidelberg. 10.1007/978-3-540-74181-7_2

Poisot, T., Canard, E., Mouillot, D., Mouquet, N., & Gravel, D. (2012). The dissimilarity of species interaction networks. Ecology Letters, 15(12), 1353–1361. 10.1111/ele.12002

Ponisio, L. C., Valdovinos, F. S., Allhoff, K. T., Gaiarsa, M. P., Barner, A., Guimarães, P. R., Hembry, D. H., Morrison, B., & Gillespie, R. (2019). A network perspective for community assembly. Frontiers in Ecology and Evolution, 7, 103. 10.3389/fevo.2019.00103

Roshni, K., Renjithkumar, C. R., Sreekanth, G. B., Raghavan, R., & Ranjeet, K. (2022). Fish community structure and functional guild composition in a large tropical estuary (Vembanad Lake, India). Environmental Science and Pollution Research, 30(11), 29635–29662. 10.1007/s11356-022-24250-8

Sandwell, D. R., Pilditch, C. A., & Lohrer, A. M. (2009). Density dependent effects of an infaunal suspension-feeding bivalve (*Austrovenus stutchburyi*) on sandflat nutrient fluxes and microphytobenthic productivity. Journal of Experimental Marine Biology and Ecology, 373(1), 16–25. 10.1016/j.jembe.2009.02.015

Schielzeth, H. (2010). Simple means to improve the interpretability of regression coefficients. Methods in Ecology and Evolution, 1(2), 103–113. 10.1111/j.2041-210X.2010.00012.x

Schleuning, M., Fründ, J., Schweiger, O., Welk, E., Albrecht, J., Albrecht, M., Beil, M., Benadi, G., Blüthgen, N., Bruelheide, H., Böhning-Gaese, K., Dehling, D. M., Dormann, C. F., Exeler, N., Farwig, N., Harpke, A., Hickler, T., Kratochwil, A., Kuhlmann, M., … Hof, C. (2016). Ecological networks are more sensitive to plant than to animal extinction under climate change. Nature Communications, 7(1), 13965. 10.1038/ncomms13965

Sekar, K. C., Thapliyal, N., Pandey, A., Joshi, B., Mukherjee, S., Bhojak, P., Bisht, M., Bhatt, D., Singh, S., & Bahukhandi, A. (2024). Plant species diversity and density patterns along altitude gradient covering high-altitude alpine regions of west himalaya, india. *Geology*, Ecology, and Landscapes, 8(4), 559–573. 10.1080/24749508.2022.2163606

Sharpe, P. J., & Baldwin, A. H. (2009). Patterns of wetland plant species richness across estuarine gradients of Chesapeake Bay. Wetlands, 29(1), 225–235. 10.1672/08-111.1

Siciliano, A., Schiel, D. R., & Thomsen, M. S. (2019). Effects of local anthropogenic stressors on a habitat cascade in an estuarine seagrass system. Marine and Freshwater Research, 70(8), 1129. 10.1071/MF18414

Simanonok, M. P., & Burkle, L. A. (2014). Partitioning interaction turnover among alpine pollination networks: Spatial, temporal, and environmental patterns. Ecosphere, 5(11), art149–art149. 10.1890/ES14-00323.1

Sponsler, D. B., Requier, F., Kallnik, K., Classen, A., Maihoff, F., Sieger, J., & Steffan-Dewenter, I. (2022). Contrasting patterns of richness, abundance, and turnover in mountain bumble bees and their floral hosts. Ecology, 103(7), e3712. 10.1002/ecy.3712

Taylor, R., & Cole, R. (1994). Mobile epifauna on subtidal brown seaweeds in northeastern New Zealand. Marine Ecology Progress Series, 115, 271–282. 10.3354/meps115271

Thomsen, M. S., Hildebrand, T., South, P. M., Foster, T., Siciliano, A., Oldach, E., & Schiel, D. R. (2016). A sixth-level habitat cascade increases biodiversity in an intertidal estuary. Ecology and Evolution, 6(22), 8291–8303. 10.1002/ece3.2499

Thomsen, M. S., & South, P. M. (2019). Communities and attachment networks associated with primary, secondary and alternative foundation species; a case study of stressed and disturbed stands of southern bull kelp. Diversity, 11(4), 56. 10.3390/d11040056

Tonkin, J. D., Bogan, M. T., Bonada, N., Rios-Touma, B., & Lytle, D. A. (2017). Seasonality and predictability shape temporal species diversity. Ecology, 98(5), 1201–1216. 10.1002/ecy.1761

Tonkin, J. D., Death, R. G., Muotka, T., Astorga, A., & Lytle, D. A. (2018). Do latitudinal gradients exist in New Zealand stream invertebrate metacommunities? PeerJ, 6, e4898. 10.7717/peerj.4898

Tylianakis, J. M., & Morris, R. J. (2017). Ecological networks across environmental gradients. Annual Review of Ecology, Evolution, and Systematics, 48(1), 25–48. 10.1146/annurev-ecolsys-110316-022821

van Dijk, J., van der Vliet, R. E., de Jong, H., Zeylmans van Emmichoven, M. J., van Hardeveld, H. A., Dekker, S. C., & Wassen, M. J. (2015). Modeling direct and indirect climate change impacts on ecological networks: A case study on breeding habitat of dutch meadow birds. Landscape Ecology, 30(5), 805–816. 10.1007/s10980-014-0140-x

Vehtari, A., Gelman, A., Simpson, D., Carpenter, B., & Bürkner, P.-C. (2021). Rank-normalization, folding, and localization: An improved *R̂* for assessing convergence of MCMC. Bayesian Analysis, 16(2). 10.1214/20-BA1221

Veríssimo, M. E. S., Medeiros, C. R., & Molozzi, J. (2024). The dispersal potential of benthic macroinvertebrates is influenced by factors acting at small spatial scales in tropical estuaries. Hydrobiologia, 851(18), 4503–4520. 10.1007/s10750-024-05603-5

Wang, T., Li, D., Tian, X., Huang, G., He, M., Wang, C., Kumbhar, A. N., & Woldemicael, A. G. (2024). Mitigating salinity stress through interactions between microalgae and different forms (free-living & alginate gel-encapsulated) of bacteria isolated from estuarine environments. Science of The Total Environment, 926, 171909. 10.1016/j.scitotenv.2024.171909

Ward, C. A., Tunney, T. D., Hale, K. R. S., O’Connor, R. F., & McCann, K. S. (2026). The rewiring of ecological networks in a variable world. Nature Reviews Biodiversity. 10.1038/s44358-026-00159-9

Wernberg, T., Thomsen, M. S., Baum, J. K., Bishop, M. J., Bruno, J. F., Coleman, M. A., Filbee-Dexter, K., Gagnon, K., He, Q., Murdiyarso, D., Rogers, K., Silliman, B. R., Smale, D. A., Starko, S., & Vanderklift, M. A. (2024). Impacts of Climate Change on Marine Foundation Species. Annual Review of Marine Science, 16(1), 247–282. 10.1146/annurev-marine-042023-093037

White, P. A., Frye, H. A., Slingsby, J. A., Silander, J. A., & Gelfand, A. E. (2024). Generative spatial generalized dissimilarity mixed modelling (SPGDMM): An enhanced approach to modelling beta diversity. Methods in Ecology and Evolution, 15(1), 214–226. 10.1111/2041-210X.14259

Woodward, G. (Ed.). (2010). Ecological networks in climate change. Elsevier.

Woolley, S. N. C., Foster, S. D., O’Hara, T. D., Wintle, B. A., & Dunstan, P. K. (2017). Characterising uncertainty in generalised dissimilarity models. Methods in Ecology and Evolution, 8(8), 985–995. 10.1111/2041-210X.12710

